# Probing complexity of microalgae mixtures with novel spectral flow cytometry approach and “virtual filtering”

**DOI:** 10.1101/516146

**Authors:** Natasha S. Barteneva, Veronika Dashkova, Ivan Vorobjev

## Abstract

Fluorescence methods are widely applied for the study of the marine and freshwater phytoplankton communities. However, identification of different microalgae populations by autofluorescent pigments remains a challenge because of the very strong signal from chlorophyll. Addressing the issue we developed a novel approach using the flexibility of spectral flow cytometry analysis (SFC) and generated a matrix of virtual filters (VF) capable to of differentiating non-chlorophyll parts of the spectrum. Using this matrix spectral emission regions of algae species were analyzed, and five major algal taxa were discriminated. These results were further applied for tracing particular microalgae taxa in the complex mixtures of laboratory and environmental algal populations. An integrated analysis of single algal events combined with unique spectral emission fingerprints and light scattering parameters of microalgae can be further used to differentiate major microalgal taxa. Our results demonstrate that spectral flow cytometer (SFC-VF) and virtual filtering approach can provide a quantitative assessing of heterogenous phytoplankton communities at single cell level spectra and be helpful in the monitoring of phytoplankton blooms.

- The research was partly presented during AQUAFLUO II Colloquium 2017, Sydney, Australia and ISAC Congress, Boston 2017

## Introduction

Phytoplankton organisms form the base of aquatic food webs and have a wide range of photosynthetic and photoprotective pigments, which are of great interest as markers to identify species in freshwater and seawater environmental samples representing different phytoplankton communities [1]. Currently, several methods are used to determine phytoplankton community structure, including microscopy, flow cytometry, spectrofluorometry, fluorescent spectroscopy, and pigments analysis by high-performance liquid chromatography (HPLC) [2]. The microscopy approach is laborious, time-consuming, and reproducibility among different research groups can be low [3]. The measurement of fluorescence spectra was extensively developed for characterizing phytoplankton taxa starting the 1970s [1, 4-9]. During the past decade, multi-channel fluorometers and scanning spectrofluorometers were applied to evaluate phytoplankton composition by measuring excitation spectra of chlorophyll ***a*** (**Chl *a***) and accessory pigments fluorescence at multiple wavelengths and creating excitation-emission matrices [10-13]. *In vivo* fluorescence methods are widely used for characterization of the phytoplankton communities, but numerous attempts to achieve a taxonomic identification of the algae taxa there remains problematic [6,9]. Measurements of fluorescence of phytoplankton communities are affected by variable biomass concentrations and therefore a varying contribution in autofluorescence signal of different microalgae subpopulations as well as inter- and intra-species pigment composition variability [14]. Till now a spectral analysis of phytoplankton communities based on spectra of averaged algal samples and can overlook a contribution of a small algal population such as cryptophytes presented in environmental samples, and also fluorescent signal might have admixture from other sources such as colored chromophoric water-dissolved organic matter and detrital pigment [15-16]. So far, any spectral approach based on averaged spectral data does not allow an actual separation of microalgae taxa contributing < 20% of the biomass in heterogeneous algal population.

The critical advantage of spectral flow cytometry (SFC) is that a measurement of complete spectrum happens from single cells with rates of hundreds and thousands of events per sec [17-19]. Moreover, SFC analysis makes possible additional differentiation of heterogeneous algal mixtures by size and granularity in the manner similar to conventional flow cytometry (FCM) [17, 20]. The emission spectrum information for every single cell can be combined with light scattering data through sequential gating on combinations of standard dot plots and histograms.

Using SFC advantages we developed a novel “virtual filtering” approach (SFC-VF) based on analysis of variable spectral emission regions in combination with light scattering-related separation of algal populations based on algae cellular size and granularity. We applied SFC-VF to differentiate and characterize different microalgae taxa in binary and multi-component mixtures as well as natural environmental microalgae assemblages and were able: (1) to distinguish of microalgal cells from phytoplankton taxa with a similar combination of pigments; and (2) to remove fluorescence signal from contaminating sources using light scatter-based gating. Moreover, differently, from FCM it makes possible separation of individual algal cells presented in heterogenous algal populations (such as cryptophytes) based on their unique spectral data.

## Methods

### Microalgae cell cultures

Microalgae cell cultures from major microalgae taxa including *Cyclotella meneghiniana* CCMP334 (diatoms), *Chlorella sp*. CCMP251 (chlorophytes), *Dinobryon divergens* CCMP3055 (chrysophytes), *Cryptomonas pyrenoidifera* CCMP1177 (cryptophytes) and *Aphanizomenon sp*. CCMP2764 (cyanobacteria) were obtained from the National Center for Marine Algae and Microbiota (Bigelow Laboratory for Ocean Sciences). Freshwater cultures *D. divergens, Aphanizomenon sp*. and *C. pyrenoidifera* were maintained in DY-V medium (modified from [20] at 14°C, 14°C and 20°C, respectively, under 150 µmoles/ m2/sec light and 12/12 L/D cycle. *Chlorella sp*. and *C. meneghiniana* were maintained in L1 medium and L1 derivative, L1-11 psi medium, respectively, at 14°C under 150 µmoles/ m2/sec light and 12/12 L/D cycle. For spectral analysis, 1000 µl volume of each culture was used to analyze single culture controls, 500 µl volume of each culture was used to analyze ten pairwise culture mixtures, and 200 µl volume of each culture to analyze a mixture of all five cultures together (ratio 1:1). Cell concentration of microalgae cultures was in the 20,000-75,000 cell mL^-1^ range.

### Environmental microalgae samples

For experiments on tracing spectral profile of cyanobacteria *Aphanizomenon sp*. CCMP2764 in environmental algal populations, samples were collected from 8 freshwater and coastal ponds in Massachusetts and Maine states, USA. Freshly collected and non-concentrated environmental samples were mixed with *Aphanizomenon sp*. culture in the following volume ratios: 100% of 2764 culture and 0% of pond sample, 50% of 2764 culture and 50% of pond sample, 10% of 2764 culture and 90% of pond sample, 5% of 2764 culture and 95% of pond sample, 1% of 2764 culture and 99% of pond sample, 0.5% of 2764 culture and 95.5% of pond sample, and 100% of pond sample. Cell concentration of collected environmental samples was in the 7,000-55,000 cell mL^-1^ range.

### Light microscopy

Images of microalgae culture cells were acquired using a confocal laser scanning microscope 780 (Zeiss, USA) and analyzed using ZEN software (Zeiss, USA) (**Fig.1A**).

### Spectral flow cytometry analysis

The spectral flow cytometer (spectral FCM) analyzer SP6800 (Sony Biotechnology Inc, USA) equipped with 488 nm, 405 nm and 638 nm lasers; 10 consecutive transparent optical prisms; and a 32-channel linear array photomultiplier (500-800 nm range for 488 nm excitation and 420-800 nm range for 405/638 lasers combination) was used for analysis of algal monocultures and environmental samples (**Fig.1B**). The instrument alignment was automatically performed using Ultra Rainbow calibration beads (Spherotech, USA) as described by Futamura et al. [22]. At least 50,000 events were collected for each sample. Environmental samples were recorded using all three available excitation sources, 488, 405 and 638 nm lasers. Single and mixed algal culture samples were recorded using blue 488 nm and violet 405 nm lasers. In order to increase non-chlorophyll based spectral differences between the algal populations, gain of PMT channels 24-30 was adjusted to 2, whereas the rest of the PMT channels were set to the maximum gain of 8. FSC gain and SSC gain were set to 17 with the threshold FSC value of 1.7% and fluorescence using Sony software v1.6 (Sony Biotechnology Inc., USA) and FlowJo software v10.2 (Treestar, USA).

### Spectral analysis of algal mixtures

Spectral data of all cells in the mixture were visualized in 488 nm laser excitation and 405 nm laser excitation spectrum charts (**Fig.1C**). Based on the most variable spectral regions, combination of virtual filters corresponding to spectrum regions in channels 15-20 (488 nm excitation) and channel 32 (488 nm excitation) (**Fig. 2C, left**), channels 31-32 (488 nm excitation) and channels V1-CH9 (405 nm excitation) (**Fig. 2C, middle**), and in channel 32 (488 nm excitation) and channels 4-15 (405 nm excitation) (**Fig. 2C, right**) were selected to achieve the best discrimination of the two cell populations.

**Figure 1:**
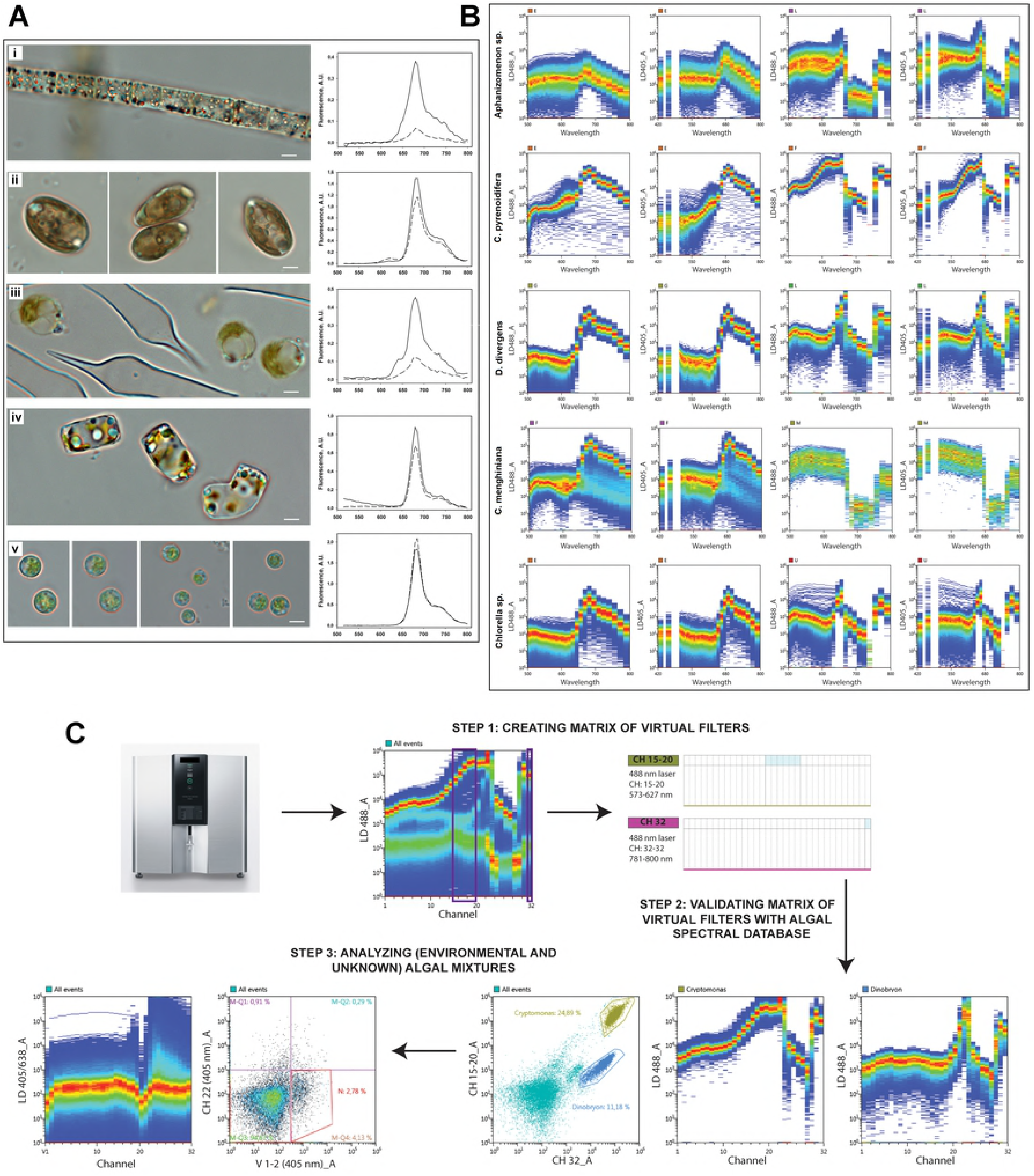
**Caption. A**. Light microscopy and spectrofluorometric data of algal cell cultures. **i** - *Aphanizomenon* sp., **ii** - *C. pyrenoidifera*, **iii** - *D. divergens*, **iv** - *C. menghiniana*, **v** - *Chlorell*a sp. First column – light microscopy images of algal cultures acquired using a confocal laser scanning microscope 780 (Carl Zeiss) in TL brightfield (objective x100); second column – spectrofluorometric data of the corresponding culture obtained with 407 nm and 488 nm excitation.**B.** Spectral FCM data of algal cell cultures. *Aphanizomenon* sp., *C. pyrenoidifera, D. divergens, C. menghiniana, Chlorell*a sp. First column – spectral data in 500-800 nm wavelength range of corresponding cultures obtained using spectral analyzer SP6800 with 488 nm laser excitation; second column – spectral data in 420-800 nm wavelength range of corresponding culture obtained using spectral analyzer SP6800 with 405 nm laser excitation; third column - spectral data in 500-800 nm wavelength range of corresponding cultures obtained using spectral analyzer SP6800 with 488 nm laser excitation and reduced intensity of channels 24-30; fourth column - spectral data in 420-800 nm wavelength range of the corresponding culture obtained using spectral analyzer SP6800 with 405 nm laser excitation and reduced intensity of channels 24-30. **C.** Virtual filtering analysis algorithm for a mixture of microalgae cells.

**Figure 2:**
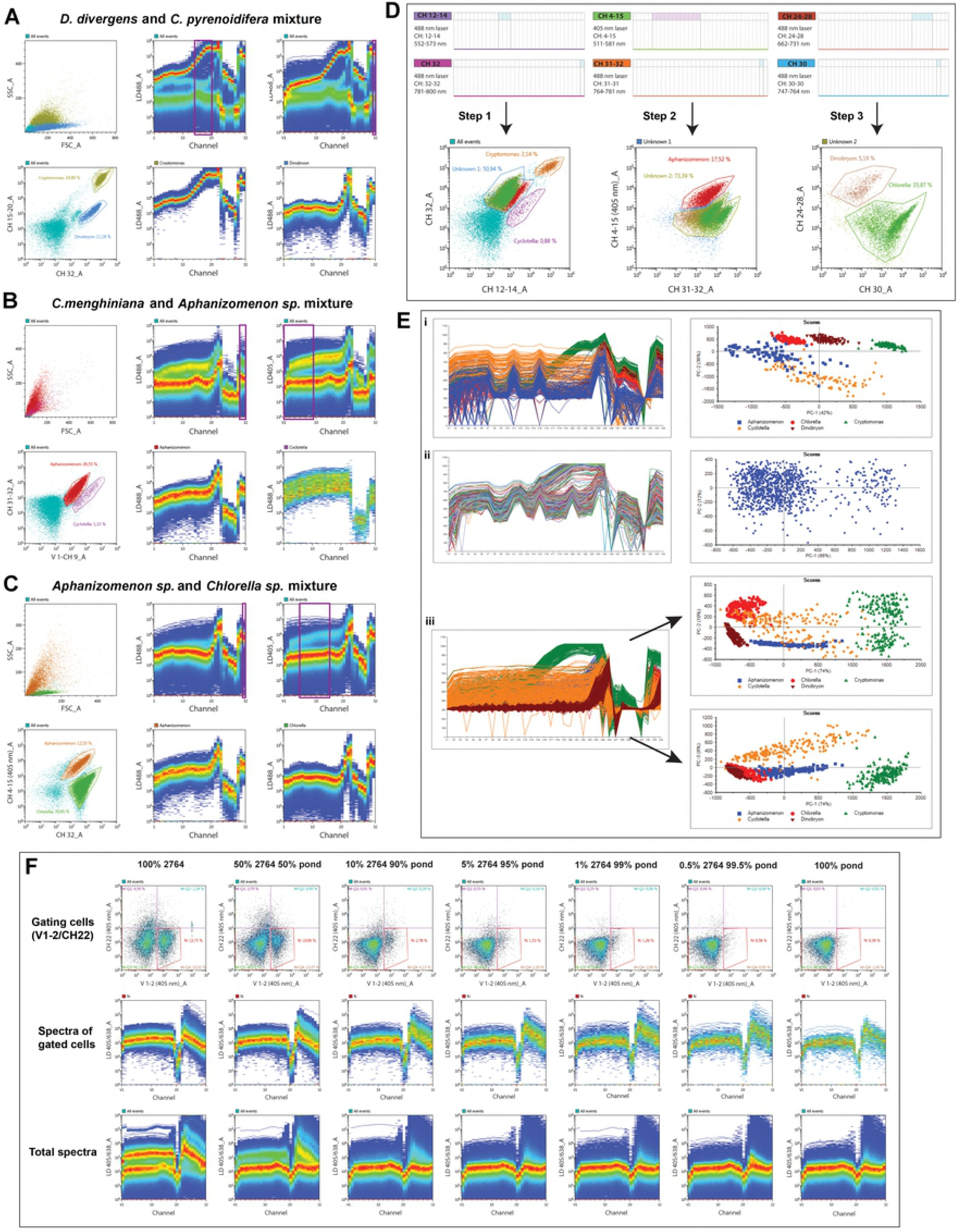
**Caption. A, B, C** Spectral analysis of double algal culture mixtures: (**A**.) *D. divergens* and *C. pyrenoidifera* spp.; (**B.**) *C. menghiniana* and *Aphanizomenon* spp. (**C.**); and *Aphanizomenon* sp. and *Chlorella* spp. **D.** Spectral analysis of five algal cultures *Aphanizomenon* sp., *C. pyrenoidifera, D. divergens, C. menghiniana* and *Chlorell*a sp. mixed together. **E.** Principal component analysis (PCA) performed for spectral data of algal cultures *Aphanizomenon* sp., *C. pyrenoidifera, D. divergens, C. menghiniana* and *Chlorell*a sp. **i** – Projection of spectra of individual cells (left) of artificially mixed algal cultures onto the plane of the first two principal components (PC) (right). **ii** - Projection of spectra of individual cells (left) of a mixture of algal cultures onto the plane of the first two PCs (right). **iii -** Projection of spectra of individual cells (left) of FCM gated populations from the mixture of algal cultures onto the plane of the PC1 and PC2 (top right) and the plane of the PC1 and PC3 (bottom right). **F.** Tracing different quantities of CCMP2764 *Aphanizomenon* sp. cells in an environmental sample from a pond based on the spectral characteristics. From left to right: 100% of 2764 cell culture, 50% volume of 2764 culture and 50% volume of pond sample, 10% volume of 2764 culture and 90% volume of pond sample, 5% volume of 2764 culture and 95% volume of pond sample, 1% volume of 2764 culture and 99% volume of pond sample, 0.5% volume of 2764 culture and 95.5% volume of pond sample, and 100% of pond sample. The first row – all cells are displayed on channel 22 (405 nm laser excitation) versus channels V1-2 (405 nm laser excitation) density plot and a region corresponding to 2764 cells region are gated (L). Second row – spectra of gated L regions are displayed on 405 nm/638 nm spectrum plots. Third row – all cells in the sample are displayed on 405 nm/638 nm spectrum plots.

Spectra of gated in specific channels populations were then plotted to confirm the identity of discriminated populations. For spectral flow cytometry analysis of five algal mixtures *C. pyrenoidifera* and *C. menghiniana* populations were separated from the mixture based on CH12-14 and CH32 (488 nm excitation) filters (**Fig. 2D, step 1**). The rest of the mixture was gated and projected onto CH4-15 (405 nm excitation) versus CH32 (488 nm excitation) dot plot to discriminate the cell population of *Aphanizomenon* sp. (**Fig. 2D, step 2**). Consequently, the unidentified population was gated and visualized on combination of CH24-28 and CH30 (488 nm excitation) filters to detach the last two populations of *D. divergens* and *Chlorella sp*. with very similar spectral profiles (**Fig. 2D**, **step 3**).

### Spectrofluorometric analysis

Spectrofluorometric data for microalgae cultures were obtained using a Varioscan Flash spectral scanning multimode reader (ThermoFisher Scientific, USA). The fluorometric scanning was performed in 515-800 nm wavelength range using 488 nm and 407 nm excitation modes. Prior to the analysis, all microalgae cultures were concentrated 30 times by centrifugation.

### Statistical analysis

Flow cytometry and spectral data were collected for at least 50,000 events for each sample and were plotted logarithmically and summarized in two-dimensional dot plots and spectrum plots. Spectral data from 500-800 nm wavelength range for 488 nm excitation (32 channel variables) and 420-800 nm wavelength range for 405 nm laser excitation (34 channel variables) were extracted from Sony software as FCS files and imported into FlowJo vs. 10.2 (Treestar, USA), where chlorophyll-positive algal populations were gated and the Area parameter of their spectra exported to comma separated values (CSV) text files. Some algal populations had a remarkably high number of cells with 0 values in channels 24-27 which may be associated with low chlorophyll signal due to dying of the cells. In order to reduce the cell heterogeneity within the sample, cells with no chlorophyll signal were removed from the population prior to the statistical analysis. The text files were then used to perform a principal component analysis (PCA) with 7 principal components using statistical software UnscramblerX v10.4 (CAMO Software, Norway). Spectral differences were also analyzed using statistical software GraphPad (GraphPad Software, USA).

### Results and Discussion

The SFC-VF method relies on identification the most variable regions of the spectra of the mixtures of algal strains analyzed pairwise, and on creating a matrix of SFC fluorescent channels corresponding to those regions. Spectral differences between single algal strains (morphology – **Fig. 1A_left_**) were captured by both spectral flow cytometer SONY SP6800 (SONY Biosciences, USA, 405 nm and 488 nm excitation) and spectrofluorometer (**Fig. 1A_right_**, **B**), however, spectrofluorometer provided an averaged signal from algal cells, debris and fluorescent organic matter. The separation of algal mixtures based on the conventional FCM approach and a filter combination used for algal analysis (such as phycoerythrin (PE) bandpass 575/25 nm) versus allophycocyanin (APC) bandpass (660/20 nm) was complicated by the heterogeneity of algal populations.

In SFC-VF approach, firstly, a sensitivity of chlorophyll-associated channels (CH24-30) captured on the SP6800 was switched to the minimal level. Then, the non-chlorophyll based spectral differences (from accessory pigments) in 420-650 nm wavelength range became prominent enabling better discrimination of algal strains (**Fig. 1C**). Further SFC analysis of algal cultures was continued with the reduced intensity of these channels.

Mixtures of algal cultures were analyzed in a pairwise manner generating ten different combinations. Initially, several variants of matrix of fluorescent channels corresponding to virtual filters capturing the algal spectra variability regions were created. We then selected a combination of fluorescent channels (virtual filter) that provides the best separation of two cell populations by dot plot. The spectra of the discriminated populations were further validated with the spectra of single algal culture controls (**Fig. 1B**). Furthermore, all five algal strains were mixed and analyzed using the spectral flow cytometry analyzer. To discriminate all algal taxa, we used a sequential gating and a combination of fluorescent channels based on virtual filters, previously selected for pairwise culture analysis (**Fig. 1C; 2A,B,C**). Consequently, the debri and fluorescent organic matter were excluded based on forward scatter/side scatter plot. Then, *C. pyrenoidifera* and *C. menghiniana* populations were separated from the other alga based on fluorescent channels CH12-14 and CH32 (488 nm excitation). The rest of the algal mixture was gated and projected onto CH4-15 (405 nm excitation) versus CH32 (488 nm excitation) dot-plot to discriminate the population of *Aphanizomenon* sp. Consequently, the initially unidentified population was gated and projected onto CH24-28 and CH30 (488 nm excitation) dot-plot to separate two populations of *D. divergens* and *Chlorella sp*. with very similar spectral profiles (**Fig. 2D**).

Spectral data recorded in 500-800 nm wavelength range for 488 nm laser excitation (32 channel variables) and 420-800 nm wavelength range for 405 nm laser excitation (34 channel variables) were used to perform a principal component analysis (PCA) of distribution of populations of algal cultures *Aphanizomenon* sp., *C. pyrenoidifera, D. divergens, C. menghiniana* and *Chlorella* sp. According to PCA results, better discrimination of algal populations was achieved using 405 nm excitation spectral data (**Fig. 2A,B,C**). PCA scores plot showed statistically significant differences between mixed algal cultures with the first principal components capturing 78% of data variation (**Fig. 2Ei**). Moreover, it allows the separation of *C. pyrenoidifera* population into two subpopulations associated with the cell heterogeneity within the culture (**Fig. 2Ei**). When algal populations are projected onto the PC1 and PC3 plane, the two *C. pyrenoidifera* subpopulations are ceased, while no discrimination of sp., *D. divergens* and *Aphanizomenon* sp. populations is observed (**Fig. 2Eiii**). However, a poor differentiation of algal populations was observed when PCA was performed on spectral data of a physical mixture of all 5 strains (**Fig. 2Eii**). Also, t-SNE cluster analysis provided less clear discrimination of microalgae subpopulations (**Supporting Fig. 1**).

**Supporting Figure 1 Capture.** Application of t-SNE analysis to the spectral data of algal cultures *Chlorell*a sp., *C. menghiniana, C. pyrenoidifera, Aphanizomenon* sp., and *D. divergens*. **A** – Color-coded file identifier columns representing FSC-H data for each strain in the merged file; **B** - t-SNE plot illustrating clusters of corresponding algal strains based on the spectral characteristics.

In the next approach we tested whether a particular microalgae type or species can be traced in the mixture of environmental microalgae populations based on its spectral profile. For this aim different quantities (from 50% to 0.5%) of *Aphanizomenon* sp. culture were mixed with environmental samples and analyzed using SFC-VF. Overall, it was possible to trace cell population of *Aphanizomenon* sp. in all eight environmental samples (an example of analysis is provided in **Fig. 2F)**. A combination of the virtual filters CH 22 (405 nm excitation) and V1-2 (405 nm excitation) enabled the best separation of *Aphanizomenon* sp. population in the 1:1 mixture of *Aphanizomenon* sp. and environmental sample (50% strain 2764 : 50% pond) and was used for analysis of other volume ratios. Single control samples of *Aphanizomenon* sp. (100% strain 2764) were used to gate the region corresponding to fluorescent live cells and compare the spectra of the gated region in different ratio mixtures. Spectra of *Aphanizomenon* sp. cells could be traced in the mixture containing as little as 0.5% proportion relative to the total volume. Notably, a small population of cells with a spectral profile similar to *Aphanizomenon* sp. was detected in the gated region of 100% pure environmental sample, which can be explained by the presence of similar or same cyanobacteria species in the collected sample.

In conventional cytometry, hardware optical filters are used to separate fluorescent signals during instrument detection. To optimize fluorescence detection and decrease acquisition of signal coming from a region with high level of autofluorescence (for example, GFP signal from cellular autofluorescence in a green-range region), would require replacement of standard optical filter with modified one [23]. In SFC software, spectral unmixing algorithms can be applied for analysis of spectral data such as “conventional” algorithm based on Least Square Method (LSM), or Weighted Least Square Method (WLSM). We applied both spectral unmixing algorithms to algal mixtures (data not shown). However, a spreading spillover from prominent **Chl *a*** led to insufficient resolution of different microalgae taxa. The SFC-VF approach [20, 24] allows the creation of “virtual bandpass filters” with no hardware modification and without spectral unmixing. As a result, it was possible to narrow or to widen spectral signal that is taken into consideration from ∼10 nm to ∼300 nm bandwidth (for SP6800 instrument) and to achieve significant discrimination of algal populations.

Here we analyzed representatives of 5 major groups of microalgae, namely (1) *Cyclotella menenginiana* from *Bacillariophyta* (diatoms); (2) *Cryptomonas pyrenoidifera* from *Cryptophyta* (cryptophytes); (3) *Aphanizomenon sp*. from *Cyanobacteria*; (4) *Chlorella sp*. from *Chlorophyta* (green algae); (5) *Dinobryon divergens* from *Ochrophyta* (chrysophytes) as model microalgal species. The data presented demonstrate the potential of our approach to the identification and quantitative evaluation of algal mixtures and experimental samples. In our study we used fresh cultures, however, there are anticipated that different preservation protocols (fixation in paraformaldehyde and freezing in liquid nitrogen) may have a smoothing effect on shape of emission spectra like it happens for absorption spectral region where absorption related to phycobilins [25]. One of main constraints in applying optical methods to phytoplankton species detection is lack of scattering data and the limited knowledge of intra-species variation in spectral emission under natural condition [25]. Previous attempts to use phytoplankton fluorescence for taxa classification were considered unsuccessful [26]. Simultaneous utilization of light scattering and excitation-emission spectral matrix results in SFC provides more accurate and consistent information that could be used for identification of major algal taxa. To quantitate abundancy of algal populations calibration beads can be used, since light scattering measurement in SFC allows for absolute counting of algal populations based on a ratio between algae and beads.

The developed novel SFC-VF approach utilizes a combination of spectral virtual filtering matrixes and light scattering and demonstrates the potential of SFC capability to distinguish fluorescence from highly overlapping autofluorescent pigments and discriminate major algal taxa (such as cryptophytes, presented in small numbers in environmental samples). The SFC-VF approach for algal taxa differentiation opens up new research areas and possibilities of algal blooms monitoring in aquatic communities.

## Acknowledgements

Authors are grateful for grant support from Ministry of Sciences, Kazakhstan to I.A.V. and N.S.B. We also thankful to Jeff Clapper and Steve Conway of Sony Biotechnology Inc., John Daley and Andrew Mason of Dana-Farber Cancer Center Flow Cytometry Core, Boston, USA for invaluable help and access to instrumentation, and to Nicole Poulton, Bigelow Laboratory of Ocean Sciences, USA for helpful advice.

